# Metabolomics and Transcriptomics unravel the mechanism of browning resistance in *Agaricus bisporus*

**DOI:** 10.1101/2021.08.13.456203

**Authors:** Zhi-Xin Cai, Mei-Yuan Chen, Yuan-Ping Lu, Zhong-Jie Guo, Zhi-Heng Zeng, Hui-Qing Zheng, Jian-Hua Liao

**Affiliations:** Institute of Edible Fungi, Fujian Academy of Agricultural Sciences, Fuzhou 350012, Fujian Province, China

## Abstract

*Agaricus bisporus* is widely used on the world market. The easy browning of mushroom surface is one of the most intuitive factors affecting consumer purchase. A certain cognition on browning mechanism have been made after years of research. At present, people slow down the browning of mushrooms mainly by improving preservation methods. In addition, breeding is also a reliable way. In the production practice, we have identified some browning resistant varieties,and we selected a browning-resistant variety to compare with ordinary variety to reveal the resistance mechanism. Using transcriptomics and metabolomics, the differences in gene expression and metabolite levels were revealed, respectively. The results showed that differentially expressed genes (DEGs) like *AbPPO4*, *AbPPO3* and *AbPPO2* were differently expressed and these DEGs involved in many pathways that related to browning. The expression of *AbPPO* expression play an important role in the browning of *A. bisporus* and multiple PPO family members are involved in the regulation of browning. However, the resistance to browning cannot be judged only by the expression level of *AbPPOs*. Formetabolomics, most of the different metabolites were organic acids. These organic acids had a higher level in anti-browning (BT) than easy-browning varieties (BS), although the profile was very heterogeneous. On the contrary, the content of trehalose in BS was significantly higher than that in BT. Higher organic acids decreased pH and further inhibited PPO activity. In addition, the BS had a higher content of trehalose, which might play roles in maintain the activity of PPO. The difference of browning resistance between BS and BT is mainly due to the differential regulation mechanism of PPO.

## Introduction

*Agaricus bisporus* (*A. bisporus*) is a kind of worldwide cultivated edible fungus with high yields and the enormous consumption. *A. bisporus* has high nutritional value with low heat and high protein. The activation of flavor amino acids and nucleotides, such as soluble sugar, polyols, free amino acids, 5’-nucleotides and sodium glutamate (MSG), in *A. bisporus* makes it a unique aroma and flavor [1, 2]. In addition to its nutritional value, *A. bisporus* is widely favored because of its numerous medicinal values. *A. bisporus* contains tyrosinase, rich selenium and vitamins, significantly reducing cholesterol and blood pressure [3]. In recent years, due to the successful realization of mushroom’s deep culture technology, people can use mushroom mycelium to produce some substances such as protein, mycosaccharide and oxalic acid. In addition, *A. bisporus* is rich in unsaturated fatty acids, which can also reduce liver cirrhosis, arteriosclerosis, obesity, heart disease and so on [4]. However, mushroom preservation has always been a complex problem in the whole industry, and browning is challenging to overcome.

Browning is one of the important factors affecting the quality of edible fungi, which seriously affected the quality and nutritional value of edible fungi. The factors of browning include internal factors (e.g., variety, moisture content, maturity, respiration rate, microorganism) and external factors (e.g., temperature, humidity, gas composition, mechanical damage, etc.). The cause of browning can be divided into two categories: enzymatic and non-enzymatic browning. Enzymatic browning is considered to be the leading cause of postharvest browning of edible fungi. Previous studies reported that the main enzymes causing browning in edible fungi are tyrosinase, such as *A. bisporus*, White Flammulina, and Coprinus [5–7]. Tyrosinase catalyzed the synthesis of L-dopa from phenolic substrates, followed by the further synthesis of dopaquinone, and finally formed melanin [8]. However, tyrosinase catalyzed phenolic substrates are different in different edible fungi. In addition, microbial infection is also the cause of the occurrence of browning of *A. bisporus*, such as *Pseudomongs tolaasii*, (*P. tolaasii*). The non-enzymatic browning mainly refers to the color deepening of mushroom surface caused by oxidation in contact with air, which might lead to the destruction of nutrients and deterioration of quality. Li et al determined the color difference, total phenol content and enzyme activity of *A. bisporus* during storage and suggested that the polyphenol oxidase (PPO) and phenylalanine ammonia lyase (PAL) in *A. bisporus* are the main factors causing browning [5].

In the present study, we collected several *A. bisporus*, including varieties with browning susceptibility, varieties with browning resistance and varieties with intermediate traits. We used transcriptional and metabolomics to analyze the browning mechanism of *A. bisporus* from different fungi. We hypothesized that browning is due to the interaction of certain genes and specific metabolites. This study will provide a theoretical basis for *A. bisporus* browning.

## Materials and methods

### Samples Preparation

Fruits of mushroom (*A. bisporus*), each with the cap arithmetic mean diameter of 30-45 mm at commercial mature stage, were harvested from local climate experiment mushroom room of the Institute of Edible Fungi, Fujian Academy of Agricultural Sciences of China, and transported under ambient conditions within 5 minutes to the laboratory. From four batches of mushroom, containing 80 fruits each, 200 samples were selected on the basis of the uniformity of maturity and appearance so that any open cap, nonwhite surface color, mechanical damaged, and defected mushrooms were excluded. In order to identify the browning sensitivity of mushrooms from different fungal sources, they were randomly divided into 3 groups (40 in each group) and stored at standard temperature and relative humidity (3±1°C and 92±2% R.H.). The fresh fruiting bodies were collected and the stipe was removed. The caps were evenly scratched with a sandpaper and stand for 10 minutes. The whiteness value of the caps surface was measured with a whiteness meter (Qtboke YQ-Z-488, Hangzhou, China). All detection were repeated three times with at least six biological and five technical replicates per treatment.

### Global gene transcriptional analysis

According to the results of browning sensitivity identification, the anti-browning (BT) and easy-browning varieties (BS) were selected for transcriptome sequencing. Two samples were randomly selected as biological repeats in each group. To construct each cDNA library, total cellular RNA was extracted from fibroblasts using Trizol reagent (Invitrogen, CA, USA). Each cDNA library was constructed according to the manufacturer’s guidelines, and next-generation sequencing was carried out in the company (Allwegene Co. Ltd, Beijing, China), using an Illumina HiSeq 4000 platform (pair-end 150bp).

For sequence data analysis, the Trimmomatic package (version 0.32) was used to obtain clean reads under default parameters. Then, the reference genome sequence of *A. bisporus* was downloaded from fungi ensemble database (http://fungi.ensembl.org/Agaricus_bisporus_var_bisporus_h97/Info/Index). All the clean reads were mapped to the reference using Tophat2 (version 2.1.1) with default parameters. After the number of reads mapped to each gene was counted, the FPKM (fragments per kilobase per million fragments) method was used for normalization through (RNA-Seq by Expectation Maximization) RSEM (https://deweylab.github.io/RSEM/), and the lowly expressed genes (FPKM < 1) were filtered in each sample [9]. DESeq2 was employed to calculate the log 2-fold change (log2FC) and probability value for each gene in every comparison, and strict criterion were used (log2FC > 2 or log2FC < −2, false discovery rate < 0.01). To analyze the potential functions of the DEGs, we enriched the DEGs according to Gene Ontology (GO) and Kyoto Encyclopedia of Genes and Genomes (KEGG) database with KOBAS (version 2.0) software. The threshold of significance was defined as FDR < 0.05.

### Sample preparation and analysis by UPLC-QTOFMS

In order to identify the difference of metabolites between the two varieties of *A. bisporus*, we selected 12 samples (six in each group, two groups) for metabolomics analysis. The untargeted metabolomics was carried out by ultra-performance liquid chromatography-quadrapole time of flight mass spectrometry (UPLC-QTOFMS) according to the previously established methods [10–12]. The freeze-dried samples were crushed using a mixer mill (MM 400, Retsch) with a zirconia bead for 1.5 min at 30 Hz. Then 100 mg powder was weighed and extracted overnight at 4°C with 1.0 ml 70% aqueous methanol containing 0.1 mg/L lidocaine for internal standard. Following centrifugation at 10000 g for 10min, the supernatant was analyzed using an LC-ESI-MS/MS system. Briefly, 2 μl of samples were injected onto a Waters ACQUITY UPLC HSS T3 C18 column (2.1 mm*100 mm, 1.8 μm) operating at 40°C and a flow rate of 0.4 mL/min. The mobile phases used were acidified water (0.04 % acetic acid) (Phase A) and acidified acetonitrile (0.04 % acetic acid) (Phase B). Compounds were separated using the following gradient: 95:5 Phase A/Phase B at 0 min; 5:95 Phase A/Phase B at 11.0 min; 5:95 Phase A/Phase B at 12.0 min; 95:5 Phase A/Phase B at 12.1 min; 95:5 Phase A/Phase B at 15.0 min. The effluent was connected to an the Synapt G2-S QTOFMS system (Waters, Milford, MA).

### Metabolomic data analysis

To produce a matrix containing fewer biased and redundant data, peaks were filtered to remove the redundant signals caused by different isotopes, in-source fragmentation, K+, Na+, and NH4+ adduct, and dimerization. According to the detected peaks, metabolites were identified by searching internal database and public databases [MassBank (https://massbank.eu/MassBank/), KNApSAcK (www.knapsackfamily.com/KNApSAcK/), HMDB (https://hmdb.ca/), MoTo DB [13], and METLIN [14]). Orthogonal projection to latent structures-discriminant analysis (OPLS-DA) was performed for classification and discriminant analysis of the samples. OPLS-DA was applied in comparison groups using R software according to a previous report [15]. A variable importance in projection (VIP) score of (O)PLS model was applied to rank the metabolites that best distinguished between two groups. The threshold of VIP was set to 1. In addition, T-test was also used as a univariate analysis for screening differential metabolites. Those with a P value of T test < 0.05 and VIP ≥ 1 were considered differential metabolites between two groups. Then, metabolites were mapped to KEGG metabolic pathways for pathway analysis and enrichment analysis.

### Statistical Analysis

Data are presented as the mean ± S.D. Statistical analyses were performed using IBM SPSS 20 (IBM, Armonk, NY). Comparison between groups was performed using one-way analysis of variance, followed by the Scheffe’s procedure for the post hoc test. P < 0.05 was considered to be statistically significant.

## Results

### Evaluation of browning resistance

The resistance identification of two varieties of BT and BS respectively, we found that BT has significantly stronger browning resistance than BS (**Figure 1A**). The whiteness value of BT exceeds 80% of the whiteness value of BS (**Figure 1B**).

**Figure 1.**
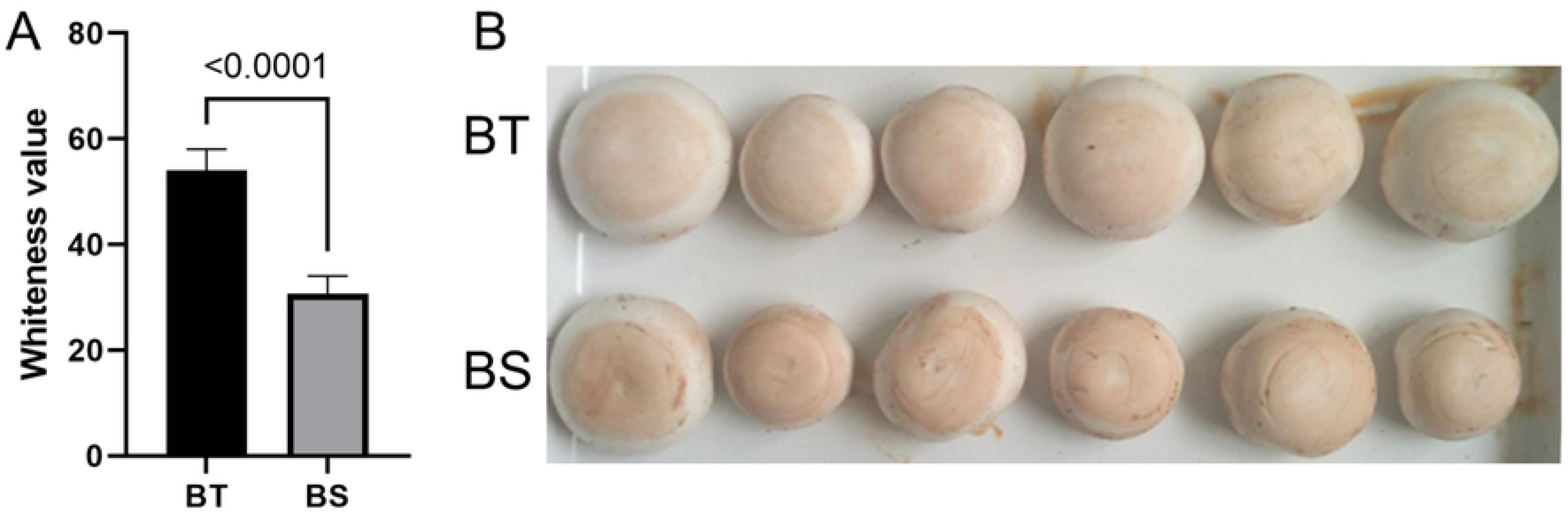
Evaluation of browning resistance BT notes the anti-browning mushroom variety; BS notes the easy-browning variety. A shows the statistical results of the whiteness values of the two varieties of mushrooms, using the T-test. B shows the the intuitive difference between the two varieties.

### Summary of transcriptome data

A total of 9,505 genes were detected in 4 samples. Percentages of reads mapped to the genome ranged from 69.27%-73.89%. After removing adapters, low-quality reads and contaminant rRNA reads, the remaining high-quality RNA-seq datasets contained 6.0-7.2 Gb of clean data from each sample (**Table S1**). The Q20, Q30 and GC content values were 96.81-97.14%, 91.87-92.46% and 49.22-49.50%, respectively (**Table S1**). Totally,1,133 DEGs were identified in the four samples, including 522 up-regulatedand 611 down-regulatedDEGs (BT *vs.* BS).

**Table S1.** Summary of transcriptome data

| Sample ID | Read Sum | Base Sum | Q20(%) | Q30(%) | GC(%) |
| --- | --- | --- | --- | --- | --- |
| BS_r1 | 24,070,648 | 7,221,194,400 | 96.94% | 92.14% | 49.39% |
| BS_r2 | 22,101,726 | 6,630,517,800 | 97.14% | 92.46% | 49.22% |
| BT_r1 | 20,196,242 | 6,058,872,600 | 96.91% | 92.03% | 49.50% |
| BT_r2 | 20,445,011 | 6,133,503,300 | 96.81% | 91.87% | 49.34% |

BT notes the anti-browning mushroom variety; BS notes the easy-browning variety. R1 and R2 means the biological replicates.

### Enrichment of functional genes

To determine the main biological functions filled by BT and BS DEGs, these transcripts were firstly mapped to GO terms in BLAST2GO analysis. The enrichment results showed that oxidation-reduction process (GO:0055114) were significantly enriched and DEGs like *AbPPO4* (upregulated), *AbPPO3* (downregulated) and *AbPPO2* (downregulated) were differently expressed.Also, melanin biosynthetic process (GO:0007093) were significantly enriched (**Figure 2A**). Interestingly, all the DEGs mapped to melanin biosynthetic process term were polyphenol oxidase members, including *AbPPO1* to *AbPPO4* and three of them were differently expressed (**Figure 2B**). The results of GO enrichment indicated that *AbPPO* gene family played a key role in mushroom browning.

**Figure 2.**
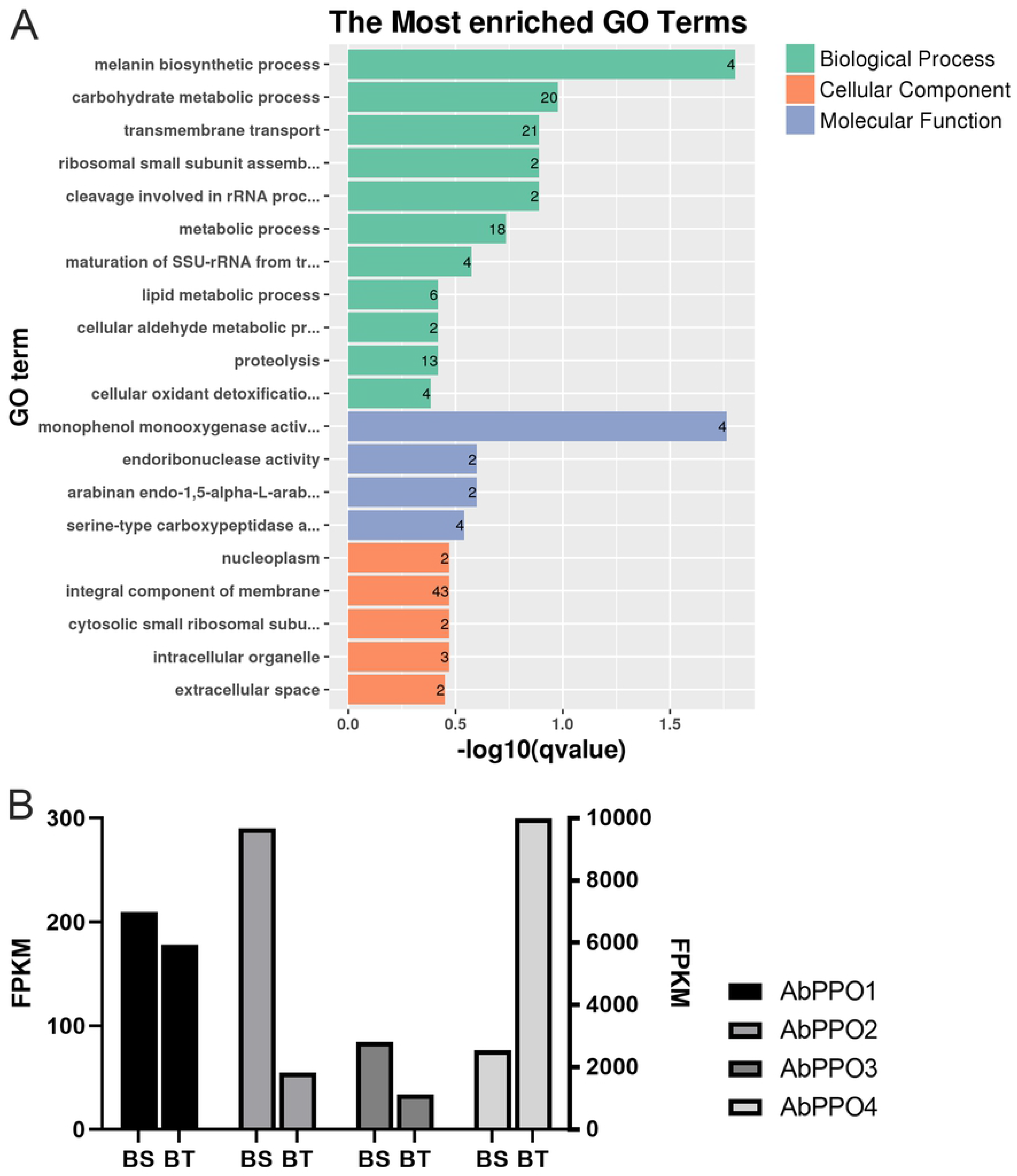
GO functional classification results. BT notes the anti-browning mushroom variety; BS notes the easy-browning variety. A shows the GO enrichment results. B shows the expression level of *AbPPO* gene family members.

Then, we performed the KEGG enrichment analysis to determine the pathways involved in DEGs. Among all the 40 significantly enriched pathways, melanogenesis (ko04916), tryptophan metabolism (ko00380), tyrosine metabolism (ko00350), and taurine and hypotaurine metabolism (ko00430) might be related to the browning of mushroom, especially the melanogenesis pathway (**Figure 3A**). Then, the expression levels of all the DEGs in these four pathways were shown in a heatmap (**Figure 3B**). We found that *PPO* family members participate in the production of melanin. Inaddition, AldA, CALM, and two hypothetical proteins (hypothetical protein AN958_09074 and hypothetical protein CVT24_011964) genes were differently expressed between BS and BT.

**Figure 3.**
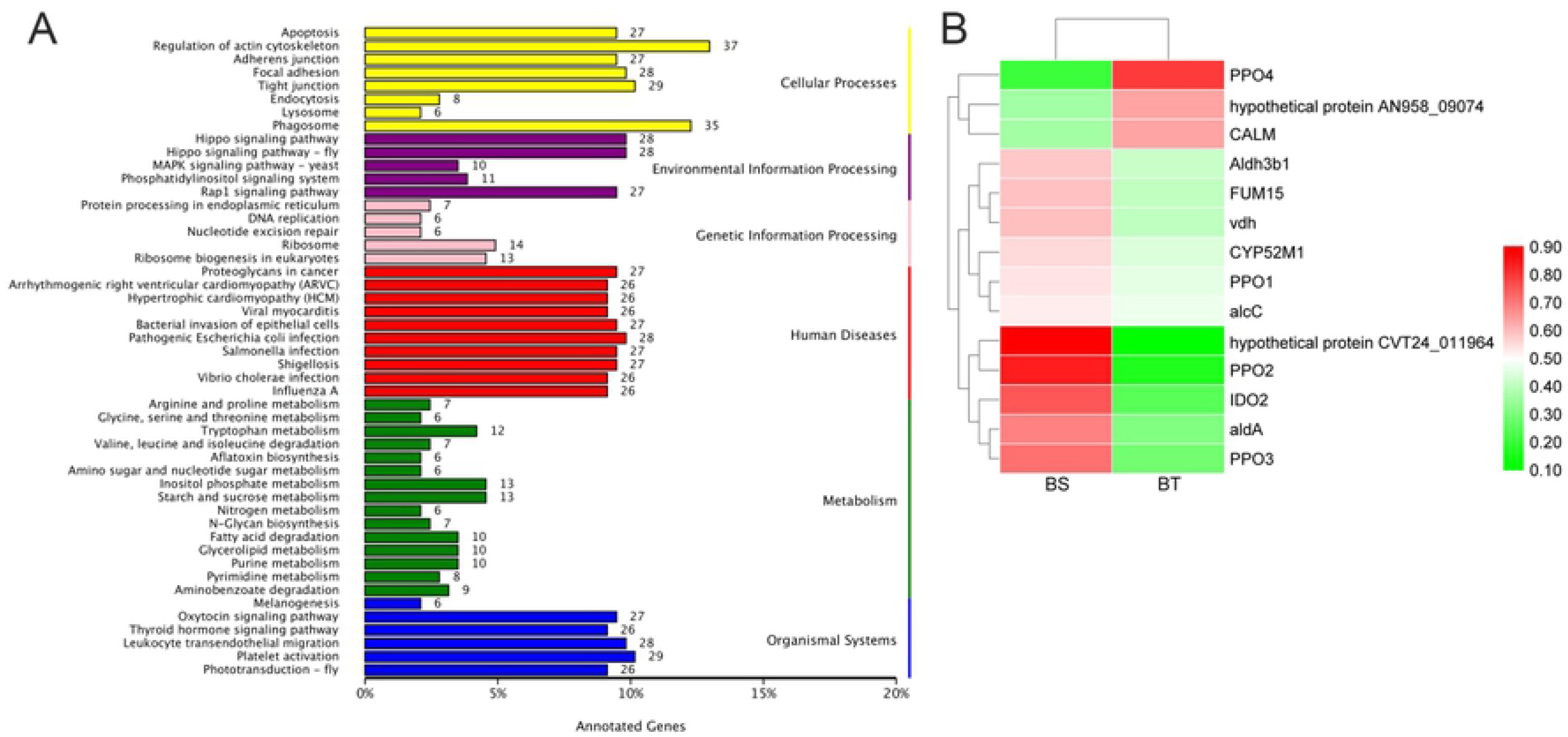
KEGG functional classification results. BT notes the anti-browning mushroom variety; BS notes the easy-browning variety. A shows the KEGG enrichment results. B shows the expression heatmap ofcandidate genes that obtained from the KEGG analysis.The red box represents the level of expression of candidate genes and the redder the color, the higher the expression level.

### Summary of metabolomic data

The metabolomics data from the present LCMS-based comparative untargeted metabolomics study were highly reproducible. OPLS-DA score plots also clearly show that there were global metabolic differences between the two groups (**Figure 4A**). Totally, we identified 474 metabolites. There were significant differences in the contents of 40 metabolites between BS and BT (including 22 metabolites without valid annotation information, **Figure 4B and C**).

**Figure 4.**
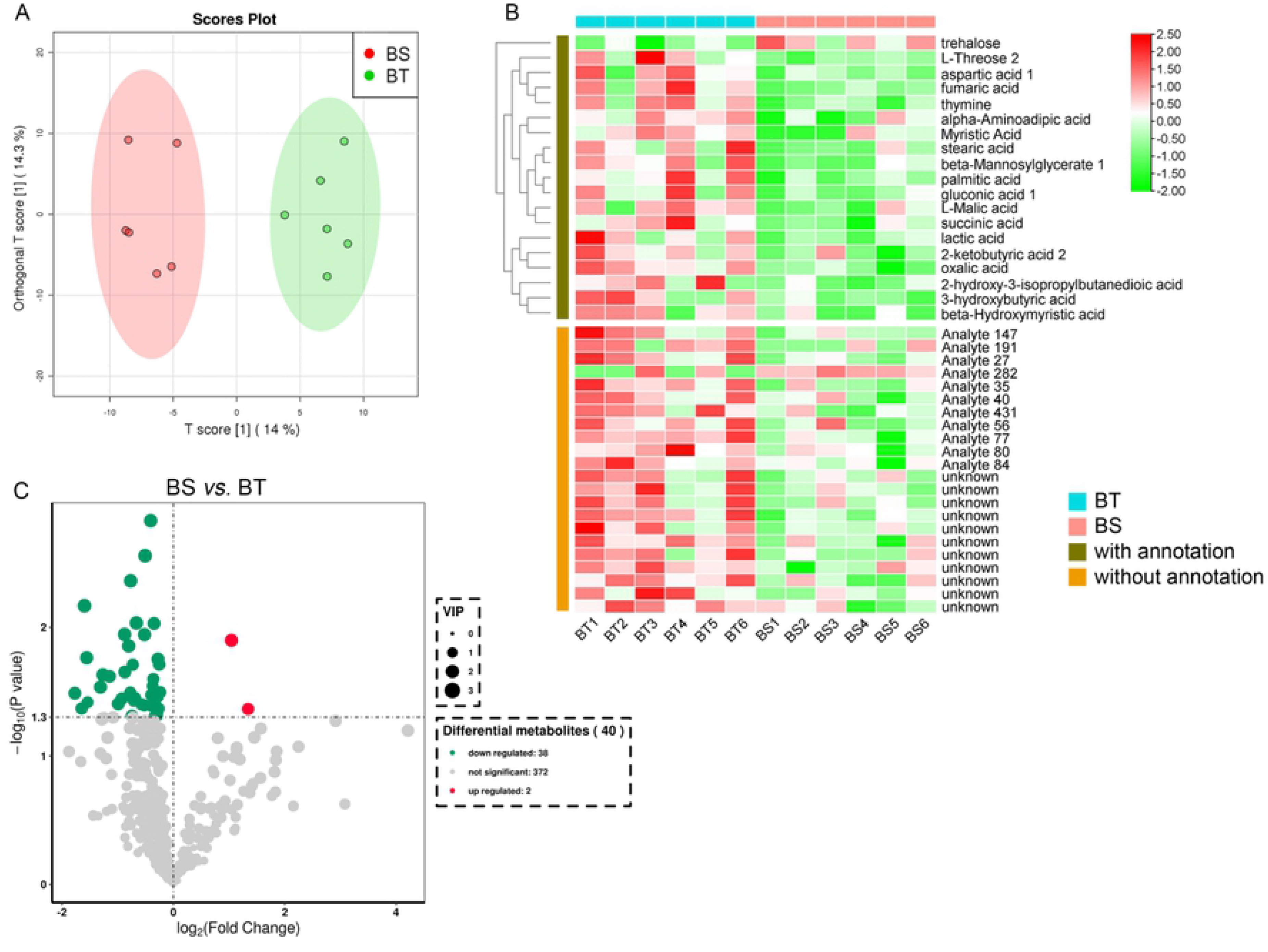
Metabonomics and differential metabolites BT notes the anti-browning mushroom variety; BS notes the easy-browning variety. A shows the PCA results of all samples. B shows the heatmap ofdifferential metabolites,the redder the color, the higher level of metabolites. C shows the volcano plot of differential metabolites.

### Differential metabolites and pathways

By annotating on all the differential metabolism, we found that most of the different metabolites were organic acids, such as succinic acid, L-Malic acid, fumaric acid, etc. These organic acids had a higher level in BT than that in BS, although the profile was very heterogeneous. On the contrary, the content of trehalose in BS was significantly higher than that in BT. In order to further analyze the function of these metabolites, we used KEGG clustering to analyze the known metabolites. We found that 15 KEGG pathways were significantly enriched, including fatty acid biosynthesis (ath00061), tyrosine metabolism (ath00350), citrate cycle (ath00020), etc (**Table 1**).

**Table 1.** KEGG enrichment results of different metabolites.

| Pathway | Hits metabolisms |
| --- | --- |
| Citrate cycle (TCA cycle) | Succinic acid;L-Malic acid;Fumaric acid |
| Fatty acid biosynthesis | Stearic acid;Myristic acid;Palmitic acid |
| Glyoxylate and dicarboxylate metabolism | L-Malic acid;Succinic acid |
| Tyrosine metabolism | Fumaric acid;Succinic acid |
| Pyruvate metabolism | L-Malic acid;L-Lactic acid |
| Alanine, aspartate and glutamate metabolism | Fumaric acid;Succinic acid |
| Biosynthesis of unsaturated fatty acids | Palmitic acid;Stearic acid |
| Fatty acid elongation in mitochondria | Palmitic acid |
| Propanoate metabolism | Succinic acid |
| Butanoate metabolism | Succinic acid |
| Carbon fixation in photosynthetic organisms | L-Malic acid |
| Glycolysis or Gluconeogenesis | L-Lactic acid |
| Starch and sucrose metabolism | Trehalose |
| Fatty acid metabolism | Palmitic acid |
| Arginine and proline metabolism | Fumaric acid |

## Discussion

In our mushroom breeding practice, we found that most of the *A. bisporus* are prone to browning, while some are resistant to browning. In order to screen the differences between the two mushroom varieties, we analyzed the transcriptome and metabolomics data of the fruiting body of the mushroom.

The transcriptome analysis showed that the *PPO* family members had the greatest effect on *A. bisporus* browning.Polyphenol oxidases are implicated in a range of biological functions in diverse systems. In addition to a role in black/brown pigment biosynthesis, PPOs may also have protective roles in plants against pathogens and environmental stress [16]. *PPO* family members participate in multiple biological pathways, and some of them proved to defense against diverse diseases and biotic stress in plants [17, 18]. As we know, in the presence of oxygen, the PPO enzyme changes substances known as phenolic compounds (through a process of oxidation) into different compounds called quinones. The quinones then react with other compounds to form melanin, then the fruit and vegetables turned brown [19, 20]. In the present results, we identified four *AbPPO* gene family members (*AbPPO1, AbPPO2, AbPPO3* and *AbPPO4*) differentially expressed between the two groups, three (*AbPPO1, AbPPO2, AbPPO3*) of which were highly expressed in BS but lower in BT. The expression of *AbPPO4* is special, whose absolute expression level was the highest of all the identified *PPO* members, and *AbPPO4* had a significantly higher expression level in BT than that in BS. These results suggest that *AbPPO1, AbPPO2, AbPPO3* and *AbPPO4* play different roles in mushroom browning, and that individual PPO gene family members are under elaborate controls to differentially express in response to specific developmental and environmental cues. According to the results of this study, higher expression of *AbPPO4* may play a positive role in the anti-browning effect of *A. bisporus*. Lei et al reported that that *AbPPO3* and *AbPPO4* contributed to the browning of mushrooms, however, the expression of *AbPPO* genes were significantly different in different parts [21]. Also, Lei found that the *AbPPO* expression trend and browning trend were different in different tissues of mushroom [21]. Hence, we speculated that the browning of *A. bisporus* is related to *AbPPO* expression, but the resistance to browning cannot be judged only by the expression level of *AbPPOs*.

Metabolomics analysis has good performance on screening the difference compounds in mushroom samples. In the present study, many organic acids (such as lactic acid, malic acid, and fumaric acid) had a higher level in BT that BS. Liu et al reported that as the concentration of citric acid increased, the activity of PPO decreased gradually [22]. Zhou reported that malic acid, citric acid and buffer with the same pH had similar effect on the relative activity of PPO, indicating the inhibition of PPO induced by malic acid and citric acid might be mainly attributed to the decrease of pH [23]. Previous studies held that acidulants could change the pH of the peripheral environment of PPO, which further induce the decrease of activity [22, 24]. Besides, organic acids inhibited PPO through chelation of the copper were reported. Yoruk and Marshall found that oxalic acid chelated the copper and diminished the catechol-quinone product formation. Kojic acid was reported to inhibit PPO by forming strong chelate with copper [25]. In this present study, L-Malic acid, succinic acid, fumaric acid, aspartic acid and palmitic acid had a higher level of organic acid. These organic acids maintained a low pH value, which inhibited the PPO activity. Additionally, the content of trehalose in all metabolites is very high, and there is a great difference between the two groups, which makes us curious about the role of trehalose in mushroom browning. Trehalose has a higher level in BS than BT. It has been shown that trehalose can protect proteins and cellular membranes from inactivation or denaturation caused by a variety of stress conditions, including desiccation, dehydration, heat, cold, and oxidation [26]. The higher content of trehalose in BS might maintained a high activity of PPO, which lead to the browning of mushroom, however, this is only our conjecture, and more experiments are needed to confirm it.

## Conclusion

The expression of *AbPPO* expression play an important role in the browning of *A. bisporus* and multiple PPO family members are involved in the regulation of browning. However, the resistance to browning cannot be judged only by the expression level of *AbPPOs*. In BT, higher organic acids decreased pH and further inhibited PPO activity. In addition, the BS had a higher content of trehalose, which might play roles in maintain the activity of PPO. Therefore, the difference of browning resistance between BS and BT is mainly due to the differential regulation mechanism of PPO.

## Acknowledgement

This research was supported by the earmarked fund of Fundamental Research Project of Fujian Provincial Research Institute for Public Welfare,China(Grant No.2019R1035-1 and 2019R1035-2), China Agriculture Research System of MOF and MARA ( CARS-20 ), the Natural Science Foundation of the Fujian Province, China(Grant No.2020J011379 ), Seed industry innovation and industrialization project of Fujian province ( zycxny2021011), and Scientific Research Project of Fujian Academy of Agricultural Sciences ( AGP2018-10 ).

## References

1. Pei F, Yang W-j, Shi Y, Sun Y, Mariga AM, Zhao L-y, Fang Y, Ma N, An X-x, Hu Q-h: Comparison of Freeze-Drying with Three Different Combinations of Drying Methods and Their Influence on Colour, Texture, Microstructure and Nutrient Retention of Button Mushroom (Agaricus bisporus) Slices. Food and Bioprocess Technology 2014, 7(3):702–710.

2. Pei F, Shi Y, Gao X, Wu F, Mariga AM, Yang W, Zhao L, An X, Xin Z, Yang F et al: Changes in non-volatile taste components of button mushroom (Agaricus bisporus) during different stages of freeze drying and freeze drying combined with microwave vacuum drying. Food Chem 2014, 165:547–554.

3. Golak-Siwulska I, Kałużewicz A, Wdowienko S, Dawidowicz L, Sobieralski K: Nutritional value and health-promoting properties of Agaricus bisporus (Lange) Imbach. Herba Polonica 2018, 64(4):71–81.

4. Chen M: The effect of different culture media on the niutrition and quality of Agaricus bisporus. Master’s thesis. Hefei: Anhui Agricultural University; 2011.

5. Nan-yi L, Qunli J, Chun-yan L, Wei-ming C: Enzymes associated with browning of Agaricus bisporus fruit bodies during storage. Acta Edulis Fungi 2009, 16:53–56.

6. Meiping Zhang YL, Yongjie Shan, Pengju Cui, Guorong Wu, Guoxiang Chen: study on the enzymatic browning mechanism and preservation of coprinus comatus. Science and Technology of Food Industry 2009, 2:84–86.

7. Lang Y: Study on Physio-Biochemical Changes Associated with Browning and Browning Mechanism of White Flammulina Velulina. Shanxi: Shanxi Agricultural University; 2014.

8. Zaidi KU, Ali AS, Ali SA, Naaz I: Microbial tyrosinases: promising enzymes for pharmaceutical, food bioprocessing, and environmental industry. Biochem Res Int 2014, 2014:854687–854687.

9. Li B, Dewey CN: RSEM: accurate transcript quantification from RNA-Seq data with or without a reference genome. BMC Bioinformatics 2011, 12:323.

10. Gustafson DL, Long ME, Bradshaw EL, Merz AL, Kerzic PJ: P450 induction alters paclitaxel pharmacokinetics and tissue distribution with multiple dosing. Cancer Chemother Pharmacol 2005, 56(3):248–254.

11. Ma X, Shah Y, Cheung C, Guo GL, Feigenbaum L, Krausz KW, Idle JR, Gonzalez FJ: The PREgnane X receptor gene-humanized mouse: a model for investigating drug-drug interactions mediated by cytochromes P450 3A. Drug Metab Dispos 2007, 35(2):194–200.

12. Lam JL, Jiang Y, Zhang T, Zhang EY, Smith BJ: Expression and functional analysis of hepatic cytochromes P450, nuclear receptors, and membrane transporters in 10- and 25-week-old db/db mice. Drug Metab Dispos 2010, 38(12):2252–2258.

13. Grennan AK: MoTo DB: a metabolic database for tomato. Plant physiology 2009, 151(4):1701–1702.

14. Zhu ZJ, Schultz AW, Wang J, Johnson CH, Yannone SM, Patti GJ, Siuzdak G: Liquid chromatography quadrupole time-of-flight mass spectrometry characterization of metabolites guided by the METLIN database. Nat Protoc 2013, 8(3):451–460.

15. Managa MG, Sultanbawa Y, Sivakumar D: Effects of Different Drying Methods on Untargeted Phenolic Metabolites, and Antioxidant Activity in Chinese Cabbage (Brassica rapa L. subsp. chinensis) and Nightshade (Solanum retroflexum Dun.). Molecules 2020, 25(6):1326.

16. Winters A, Heywood S, Farrar K, Donnison I, Thomas A, Webb KJ: Identification of an extensive gene cluster among a family of PPOs in Trifolium pratense L. (red clover) using a large insert BAC library. BMC plant biology 2009, 9:94–94.

17. Li L, Steffens JC: Overexpression of polyphenol oxidase in transgenic tomato plants results in enhanced bacterial disease resistance. Planta 2002, 215(2):239–247.

18. Duan C, Yu J, Bai J, Zhu Z, Wang X: Induced defense responses in rice plants against small brown planthopper infestation. The Crop Journal 2014, 2(1):55–62.

19. Tang H: Regulation and function of the melanization reaction in Drosophila. Fly (Austin) 2009, 3(1):105–111.

20. Twort VG, Newcomb RD, Buckley TR: New Zealand Tree and Giant Wētā (Orthoptera) Transcriptomics Reveal Divergent Selection Patterns in Metabolic Loci. Genome Biol Evol 2019, 11(4):1293–1306.

21. Lei J, Li B, Zhang N, Yan R, Guan W, Brennan CS, Gao H, Peng B: Effects of UV-C treatment on browning and the expression of polyphenol oxidase (PPO) genes in different tissues of Agaricus bisporus during cold storage. Postharvest Biology and Technology 2018, 139:99–105.

22. Liu W, Zou L-q, Liu J-p, Zhang Z-q, Liu C-m, Liang R-h: The effect of citric acid on the activity, thermodynamics and conformation of mushroom polyphenoloxidase. Food Chemistry 2013, 140(1):289–295.

23. Zhou L, Liu W, Xiong Z, Zou L, Chen J, Liu J, Zhong J: Different modes of inhibition for organic acids on polyphenoloxidase. Food Chemistry 2016, 199:439–446.

24. Queiroz C, da Silva AJR, Lopes MLM, Fialho E, Valente-Mesquita VL: Polyphenol oxidase activity, phenolic acid composition and browning in cashew apple (Anacardium occidentale, L.) after processing. Food Chemistry 2011, 125(1):128–132.

25. Battaini G, Monzani E, Casella L, Santagostini L, Pagliarin R: Inhibition of the catecholase activity of biomimetic dinuclear copper complexes by kojic acid. JBIC Journal of Biological Inorganic Chemistry 2000, 5(2):262–268.

26. Elbein AD, Pan YT, Pastuszak I, Carroll D: New insights on trehalose: a multifunctional molecule. Glycobiology 2003, 13(4):17R–27R.

